# Anti-inflammatory Activity of Lefamulin in a Lipopolysaccharide-Induced Lung Neutrophilia Model

**DOI:** 10.1101/2020.06.23.168393

**Authors:** Michael Hafner, Susanne Paukner, Wolfgang W. Wicha, Boška Hrvačić, Steven P. Gelone

**Author notes:** Correspondence to*: Steven P. Gelone, PharmD.

## Abstract

Lefamulin is a novel pleuromutilin antibiotic approved for the treatment of community-acquired bacterial pneumonia. This study demonstrated anti-inflammatory activity of lefamulin in a murine lipopolysaccharide-induced lung neutrophilia model. Pretreatment of mice at clinically relevant lefamulin subcutaneous doses (35, 70, 140 mg/kg [free base]) followed by intranasal lipopolysaccharide challenge (5 μg/50 μL/mouse) demonstrated significant, dose-dependent reductions in total and neutrophil cell counts in bronchoalveolar lavage fluid samples, with reductions comparable to oral dexamethasone (0.5 mg/kg) pretreatment.

---

Anti-inflammatory and immunomodulatory activities have been observed with various antibiotics, including macrolides, tetracyclines, and sulfonamides, which has resulted in their use in the treatment of chronic inflammatory pulmonary disorders (1). Acute lung injury (ALI) and its most severe form, acute respiratory distress syndrome (ARDS), cause inflammatory damage to the alveolar capillary membrane and excessive uncontrolled pulmonary inflammation (2). ALI/ARDS complicate pneumonia and contribute substantially to morbidity and mortality in these patients (2–4). Lipopolysaccharide (LPS) administration can induce pathologic and biological changes similar to those seen in ARDS, including neutrophilic infiltration and increased intrapulmonary cytokines, which have been extensively studied in ALI experimental models (5). Further, research has suggested that increased neutrophil recruitment to the lungs may contribute to tissue damage, particularly in chronic diseases (6).

Pleuromutilin antibiotics inhibit bacterial protein synthesis by binding to the peptidyl transferase center of the 50S ribosomal subunit (7), and lefamulin is the first pleuromutilin antibiotic approved for intravenous and oral use in humans (8). Lefamulin has demonstrated potent *in vitro* activity against the pathogens that most commonly cause community-acquired bacterial pneumonia (CABP) (9–12) and based on the results of two phase 3 clinical trials (13,14), is approved in the United States and has received a positive opinion from the Committee for Medicinal Products for Human Use in the EU for the treatment of adults with CABP (8). The current study evaluated the anti-inflammatory effects of lefamulin in a murine LPS-induced lung neutrophilia model.

## Animals

Six-week-old male BALB/c mice (Charles River, Calco, Italy) were singly housed in a temperature-controlled (22°C ± 2°C) environment with a 12:12-hour light:dark cycle and free access to food and water. Mice were given ≥7 days for acclimation before all procedures. One day before the start of experimental procedures, all animals were randomized into six groups (*n*=8/group). All animal experiments were conducted according to European Union directive 2010/63/EU and national legislation (Official Gazette 55/13) regulating use of laboratory animals in scientific research, with oversight from an Institutional Committee on Animal Research Ethics (CARE-Zg).

## Reagents

LPS lyophilized powder (2 mg; Sigma, Munich, Germany) from *Escherichia coli* (O111:B4) was dissolved in 10 mL cold saline (0.9%), vortexed, and further diluted by mixing 2 mL of this solution with 2 mL saline to reach a final concentration of 5 μg LPS/50 μL saline. Dexamethasone (Sigma) was dissolved in carboxymethylcellulose (0.5% in water) and dosed in a volume of 10 mL/kg per mouse. Lefamulin (BC-3781.Ac; Nabriva Therapeutics, Vienna, Austria) was weighed using a correction factor of 1.12 and dissolved in saline. For the three dose groups (free base: 35, 70, and 140 mg/kg), corresponding lefamulin concentrations were 3.92, 7.84, and 15.68 mg/mL (free base: 3.5, 7.0, and 14.0 mg/mL), respectively. Lefamulin was dosed in a volume of 10 mL/kg per mouse. Ketamine hydrochloride (Narketan 10) was acquired from Vetoquinol (Bern, Switzerland) and xylazine hydrochloride (Rompun, 2%) was acquired from Bayer (Leverkusen, Germany).

## Induction of Lung Neutrophilia and Treatments

Dexamethasone (0.5 mg/kg) was administered orally 1 hour before the LPS challenge. Vehicle and lefamulin were administered subcutaneously (SC) 30 minutes before the LPS challenge. Immediately before the LPS challenge, mice were anesthetized via intraperitoneal (IP) injection of ketamine (2 mg/mouse) and xylazine (0.08 mg/mouse). To induce pulmonary neutrophilia, mice in the control group received 50 μL saline and all other animals received 5 μg LPS/50 μL saline intranasally.

## Bronchoalveolar Lavage Fluid Collection and Analysis

Approximately 4 hours after LPS administration, mice were euthanized by an overdose of IP ketamine (200 mg/kg) and xylazine (16 mg/kg). Tracheostomy was performed and a Buster cat catheter (1.0 × 130 mm; Kruuse, Langeskov, Denmark), shortened to 3 cm, was clamped into the trachea. The lungs were washed three times with cold phosphate buffered saline (PBS) in a total volume of 1 mL (0.4, 0.3, and 0.3 mL). The collected bronchoalveolar lavage fluid (BALF) samples were centrifuged at 3500 rpm for 5 minutes at 4°C, and resulting cell pellets were each resuspended in 600 μL PBS. Samples were immediately analyzed for total and differential neutrophil cell counts via automated hematology analyzer (XT-2000iV; Sysmex, Kobe, Japan).

## Statistical Analyses

Statistical analyses were performed using GraphPad Prism v5.04 (GraphPad Software, Inc., La Jolla, CA, USA). Between-group differences were determined using the Mann-Whitney test and were considered statistically significant when *P*<0.05.

To our knowledge, this is the first study to investigate the anti-inflammatory activity of lefamulin. In this mouse model of lung neutrophilia, pretreatment with lefamulin at doses of 35, 70, and 140 mg/kg SC 30 minutes before intranasal LPS challenge was associated with almost complete reduction in total cell and neutrophil recruitment to the lungs at 4 hours post challenge compared with the vehicle control group (no treatment, **Figure**). Compared with dexamethasone (0.5 mg/kg), a known potent anti-inflammatory glucocorticoid, lefamulin demonstrated similar reductions in BALF total cell and neutrophil counts, even at the lowest dose (35 mg/kg). Inhibition of neutrophilic lung infiltration may be beneficial during the early phase of ARDS, which is characterized by disruption of alveolar epithelial and endothelial barriers as well as widespread neutrophilic alveolitis, leading to formation of protein-rich edema in interstitium and alveolar spaces (2,15).

Maximum plasma concentrations of 1.7 and 2.5 mg/L have been achieved in healthy mice treated with single 35 and 70 mg/kg SC lefamulin doses, respectively, and a concentration of 2.5 mg/L was achieved in healthy subjects treated with a single 150 mg intravenous lefamulin dose (16,17). Because this corresponds to the approved clinical dose (150 mg intravenous/600 mg oral) in patients with CABP (8,18), these data indicate that lefamulin anti-inflammatory activity is seen with a plasma exposure that is lower than the corresponding antimicrobial clinical dose.

In earlier investigations, the veterinary pleuromutilin antibiotic valnemulin showed *in vitro* and *in vivo* anti-inflammatory effects (19,20). LPS-induced pulmonary edema, accumulation of inflammatory cells in BALF (eg, neutrophils and macrophages), and increased inflammatory cytokines (eg, tumor necrosis factor-alpha [TNFα] and interleukin-6 [IL-6]) were significantly attenuated in mice pretreated with valnemulin or dexamethasone compared with no-treatment control group, with histologic analysis suggesting a protective effect from valnemulin on LPS-induced ALI (19). Valnemulin treatment in murine macrophages also significantly inhibited LPS-induced production of inflammatory mediators, including nitric oxide, prostaglandin E_2_, TNF-α, and IL-6 (20). Likewise, significantly reduced TNFα, IL-6, and monocyte chemoattractant protein-1 serum levels were observed in a methicillin-resistant *Staphylococcus aureus* wound infection mouse model following treatment with amphenmulin, a pleuromutilin derivative currently in development for veterinary use (21); due to the model used, however, further studies are needed to determine if these effects were the direct result of anti-inflammatory activity, an indirect consequence of antimicrobial activity, or both.

Taken together with the data presented herein, these results suggest that lefamulin has anti-inflammatory properties similar to those of macrolide antibiotics, which are currently used as anti-inflammatory therapy for pulmonary disorders (22). Like macrolides, lefamulin inhibits bacterial protein synthesis and demonstrates excellent tissue penetration, accumulation in macrophages, and immunomodulatory effects (eg, macrophage activation, neutrophilic inflammation inhibition) (8,16,22,23). Moreover, azithromycin or clarithromycin pretreatment in LPS-induced ALI similarly resulted in significantly reduced neutrophil recruitment (24). Further research on the anti-inflammatory and immunomodulatory properties of lefamulin and its potential as a treatment for inflammatory lung diseases is warranted.

## Acknowledgments

This research was funded by Nabriva Therapeutics, King of Prussia, PA, USA. Editorial and medical writing support for manuscript development was provided by Lauriaselle Afanador, PhD, Michael S. McNamara, MS, and Morgan C. Hill, PhD, employees of ICON (North Wales, PA, USA), and funded by Nabriva Therapeutics.

---

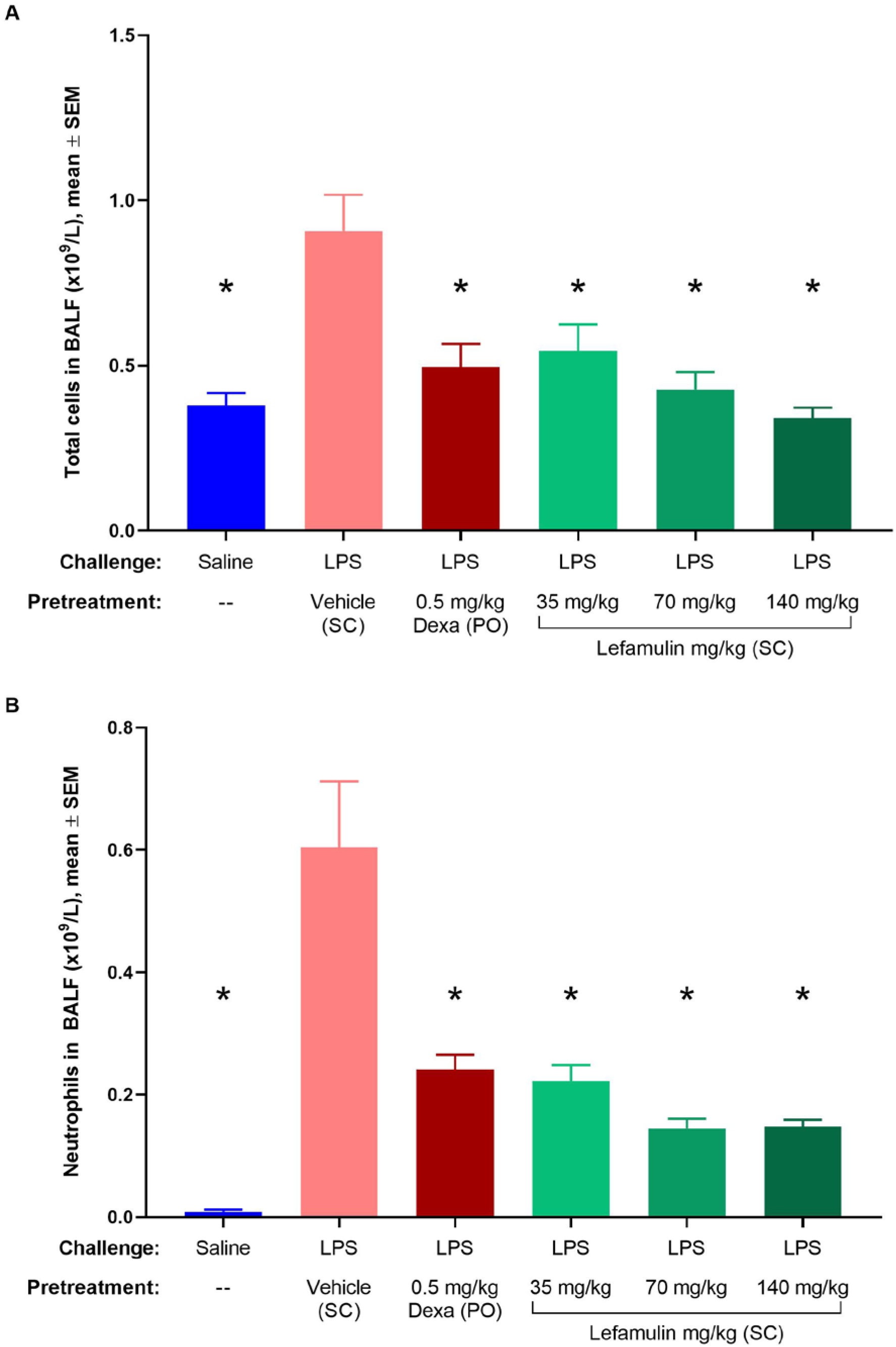
(A) Total and (B) neutrophil cell counts in BALF after LPS induction. BALF=bronchoalveolar lavage fluid; Dexa=dexamethasone; LPS=lipopolysaccharide; PO=oral; SC=subcutaneous; SEM=standard error of the mean. \**P*<0.05 vs LPS/vehicle via Mann-Whitney test.

